## Supplemental File for "Neural synchronization is strongest to the spectral flux of slow music and depends on familiarity and beat salience"

### Figure Supplements

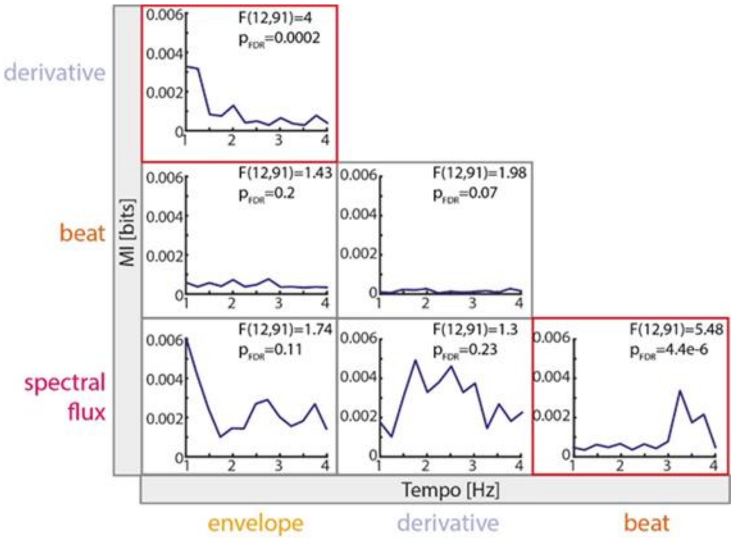


**Figure 1 – figure supplement 1. Shared Mutual Information (MI) between musical features across tempo conditions.** MI scores for all possible feature combinations as a function of tempo. Significance was evaluated based on a three-way ANOVA (with tempo as second factor, see *Material and Methods* for further information) and a follow-up pairwise comparison. Red boxes indicate significant tempo-dependent shared MI between musical features with a significance level of $p_{FDR}<0.05$.

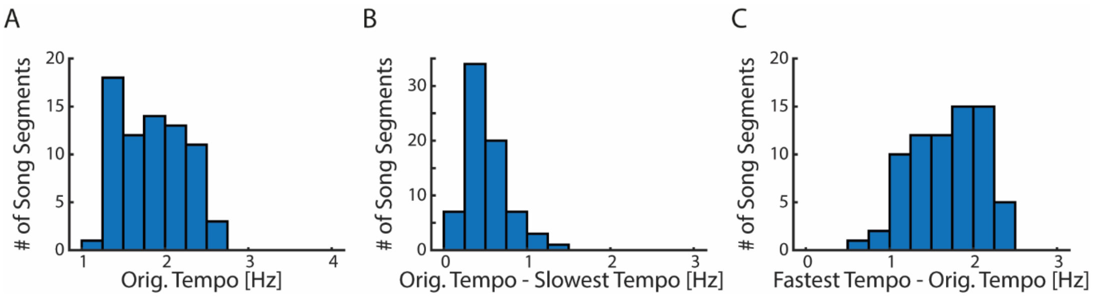


**Figure 1 – figure supplement 2. Tempo manipulations of original music segments. (A)** Histogram of the original tempo of the music segments prior to tempo manipulation binned between 1-4 Hz in steps of 0.25 Hz (n=72). Histograms of **(B)** the difference between the original tempo and the slowest possible tempo manipulation and **(C)** the difference between the fastest possible tempo manipulation and the original tempo per music segment.

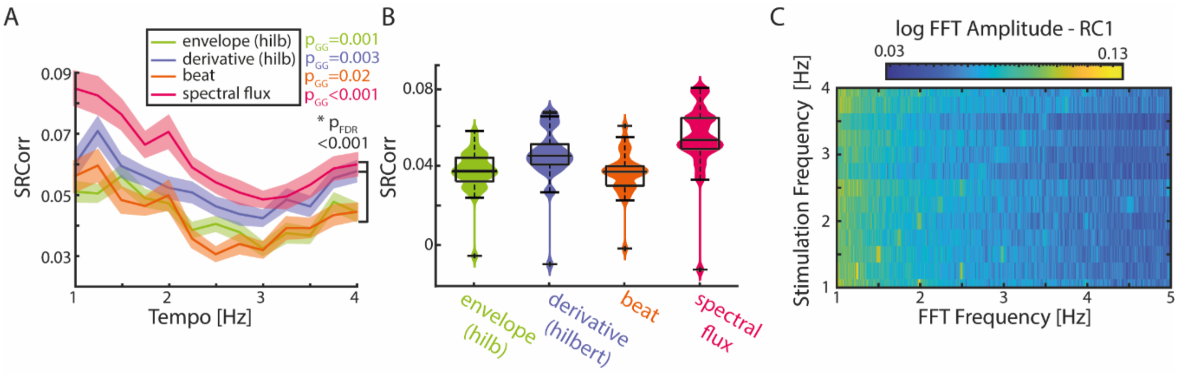


**Figure 2 – figure supplement 1. SRCorr and SRCoh in response to the full-band amplitude envelope and derivative. (A)** Mean SRCorr across stimulation tempi and musical features (± SEM). Similar to Figure 2B of the main manuscript, the full-band amplitude envelope (Hilbert transform) and resultant first derivative were used. Significance between tempi was assessed using a repeated-measure ANOVA (with Greenhouse-Geiser correction if applicable). **(B)** SRCorr across musical features. Statistically significant differences were identified between all musical feature combinations except between the envelope and beat onsets using a repeated-measure ANOVA ($p_{FDR}$<0.001). **(C)** Colormap of the fast Fourier Transform (FFT) of the first reliable component (RC1) across stimulation tempi. Note that the colorbar is in a logarithmic scale.

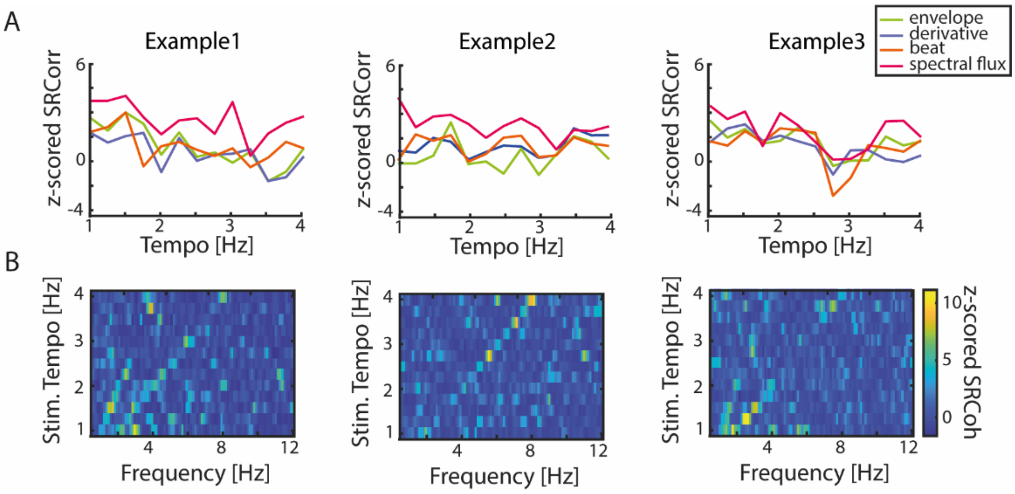


**Figure 2 – figure supplement 2. Individual data examples for the SRCorr and SRCoh. (A)** Each plot shows the mean z-scored SRCorr of one participant across stimulation tempi. Each line represents the average of one musical feature (±SEM). **(B)** Illustrative color plots of the normalized SRCoh in response to the spectral flux across stimulation tempi (1-4 Hz) of the same four participants as in (A). Z-scoring was based on the surrogate distribution.

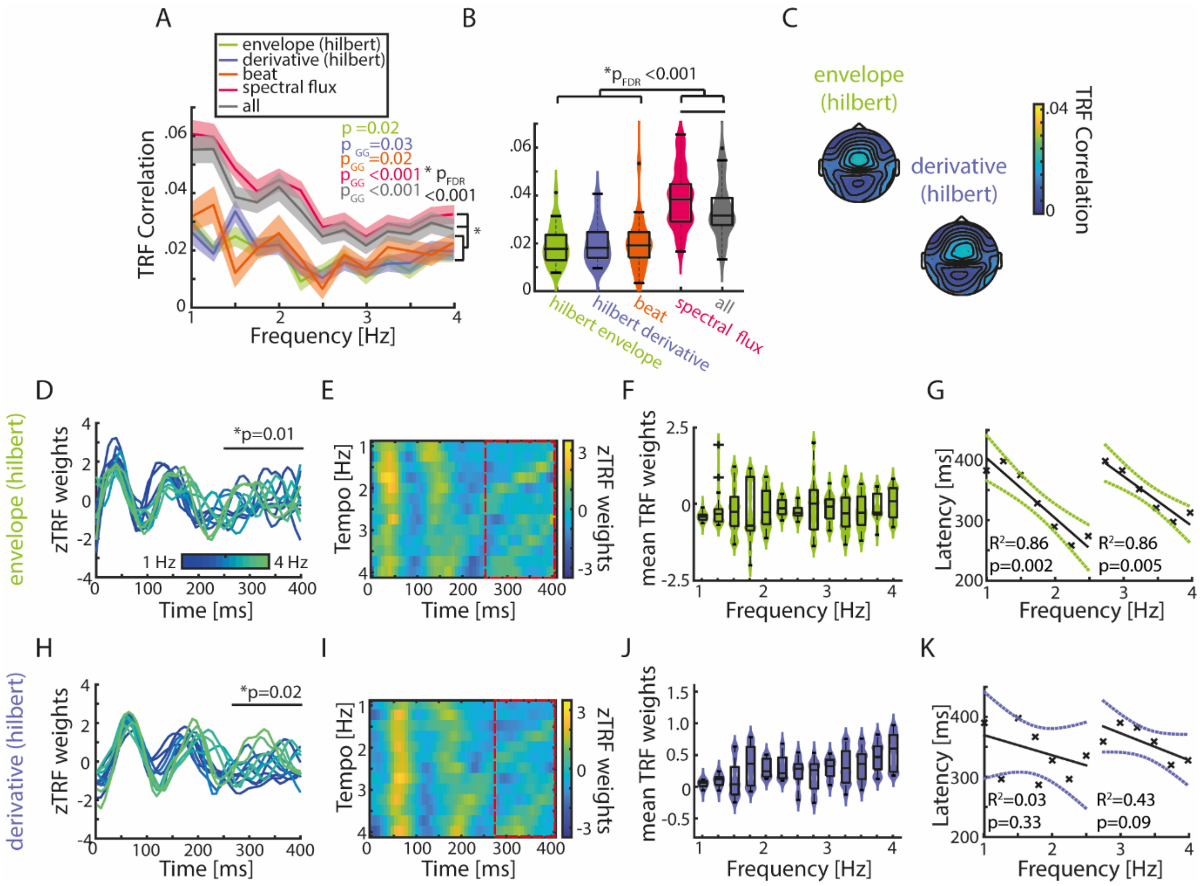


**Figure 3 – figure supplement 1. TRFs in response to the full-band amplitude envelope and first derivative show similar patterns as the gammatone filtered musical features.** **(A)** Mean TRF correlations across stimulation tempi and musical features (±SEM). **(B)** TRF correlations across musical features. Violin plots indicate the median, 25$^{th}$ and 75$^{th}$ percentiles (repeated-measure ANOVA; *$p_{FDR}$<0.001). Similarly to the gammatone filtered features, the full-band envelope and derivative show significantly smaller TRF correlations in comparison to the spectral flux and the combination of all features. **(C)** Auditory topographies of the TRF correlations in response to the full-band features. **(D)** TRF weights for time lags between 0 and 400 ms of the Hilbert envelope. Cluster-based permutation testing was used to identify significant time lag windows (envelope: 250-400 ms, p=0.01; derivative: 281-400 ms, p=0.02). Each line represents one stimulation tempo (n=13, blue, 1 Hz - green, 4 Hz). **(E)** Colormaps of the TRF weights over the same time lags for the envelope. Red dashed lines highlight significant time windows. **(F)** Average TRF weights from the significant time lag window for the envelope. **(G)** Latency of the P3 peak located in the significant time lag window across stimulation tempi for the full-band envelope TRFs. The latencies were divided into two subgroups (1-2.5 Hz and 2.75-4 Hz) and a regression fit into the data. **(H)-(K)** Same as (D)-(G) for the first derivative.

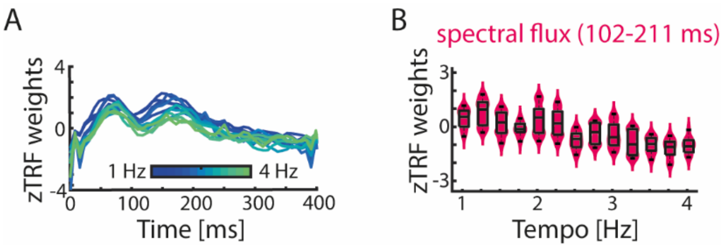


**Figure 3 – figure supplement 2. "Corrected" TRF weights of the spectral flux after removing the effects of the other musical features. (A)** TRF weights in response to the spectral flux. To calculate those weights, a multivariate TRF approach based on the amplitude envelope, first derivative and beat onsets was used and the resulting TRF predictions were subtracted from the "actual" EEG data. The residual EEG data was used to compute the spectral TRF model. **(B)** Similarly to the main Figure 3F the TRF weights at the previously calculated significant time window were plotted as a function of tempo.

26

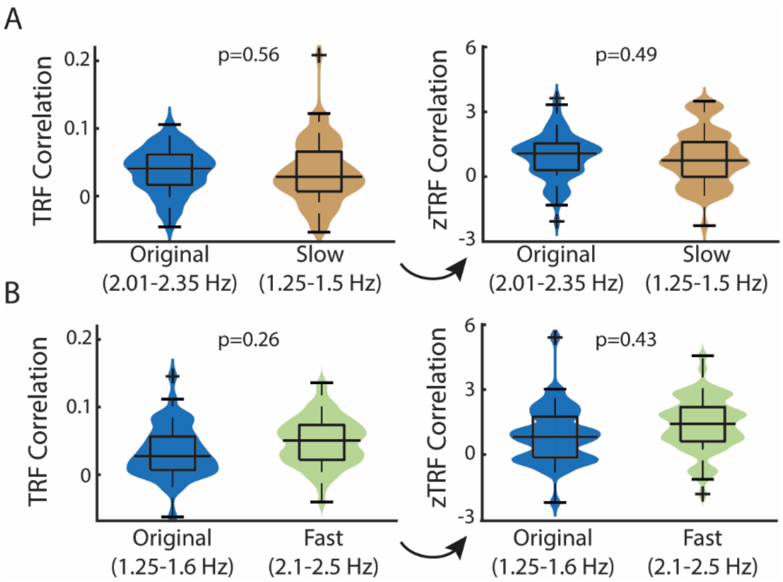


**Figure 3 – figure supplement 3. No differences in TRFs correlations between more vs. less modulated music. (A)** TRF correlations for up to three trials per participant when the original music tempo ≈ manipulated music tempo (labelled "Original") vs. when the manipulated music tempo was faster than the original music segments ("Slow"). For this analysis different trials for the different stimulation subgroups from the same stimulation tempo condition (here: 2.25 Hz) were used ($N_{ori}$=91; $N_{slow}$=96 trials). In the right plot the TRF correlation were z-scored based on the surrogate distribution on a per trials basis. No significant differences were observed between groups (repeated-measures ANOVA). **(B)** Same as (A), but here the original tempo was at a slower tempo and was contrasted against music segments that were originally faster and were manipulated to be played at a 1.5 Hz ($N_{ori}$=57; $N_{fast}$=95 trials).

27

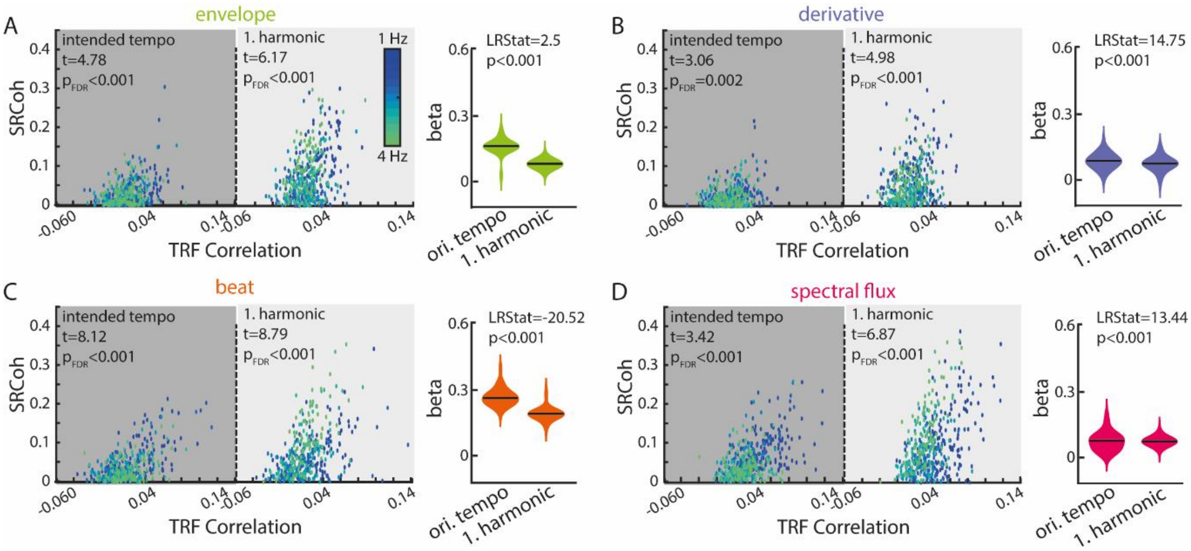


**Figure 4 – figure supplement 1. Significant relationships between SRCoh and TRF correlations for all musical features at the stimulation tempo and first harmonic. (A)** Linear-mixed effects models of the SRCoh (predictor variable) and TRF correlations (response variable) in response to the amplitude envelope at the intended stimulation tempo (left, dark grey) and first harmonic (right, light grey). Each dot represents the mean correlation of one participant (N=34) at one stimulation tempo (n=13) (=grouping variables; blue, 1 Hz-green, 4 Hz). Violin plots illustrate fixed effects coefficients (β). **(B)-(D)** same as (A) for the first derivative, beat onsets and spectral flux. For all musical features, the fixed effects were significant ($p_{FDR}$<0.01). Model comparisons were implemented based in the Likelihood ratio test (LRStat, p<0.001).

28

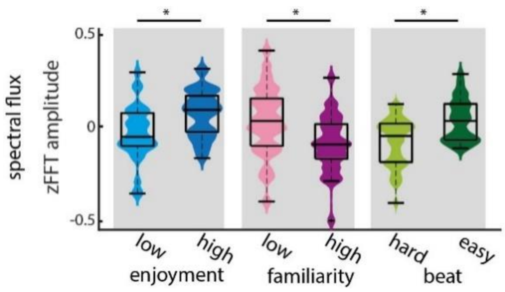


**Figure 5 – figure supplement 1. Significant differences of FFT amplitudes at stimulus-relevant frequencies between differently rated trials.** Z-scored average FFT amplitudes at the stimulation tempo and first harmonic of the 15 highest vs. lowest ratings per behavioral rating category (N=34, low vs. highly enjoyed, low vs. highly familiar, and subjectively difficult vs. easy beat trials). Significant differences were observed for all pairwise comparisons (paired-sample t-test, $p_{FDR}$≈0.01).

29

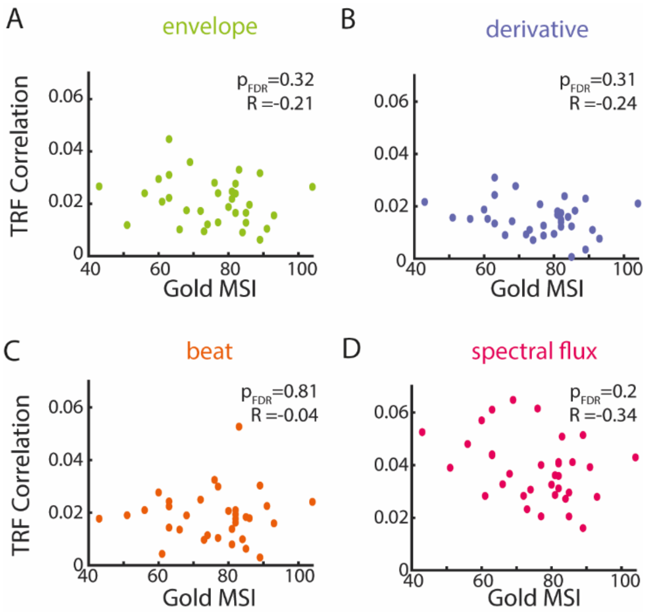


**Figure 5 – figure supplement 2. Musical training did not have an effect on TRF correlations regardless of the musical feature.** Scatter plot between the general sophistication index (F7, Gold-MSI) and mean TRF correlations per participant (n=34). No significant correlations between the Gold-MSI and TRF correlations were observed in response to the **(A)** amplitude envelope, **(B)** first derivative, **(C)** beat onsets and **(D)** spectral flux.

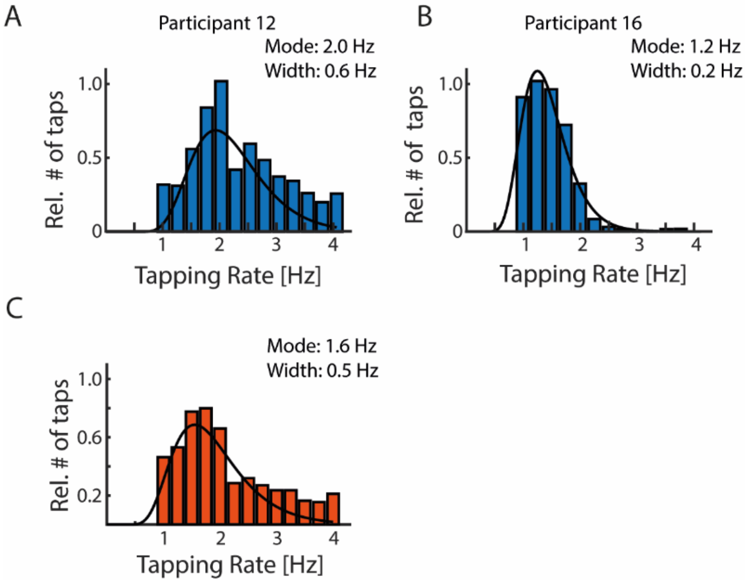


**Figure 5 – figure supplement 3. Music tapping rate across participants. (A)-(B)** Illustrative histograms of the relative number of trials per tapped music rate of two participants with a fitted skewed Gaussian. The modes indicate the preferred music tapping rate and the width the shape of the fitted Gaussian. **(C)** Mean music-tapping histogram across all participants (n=29, 5 participants were excluded from the music tapping analysis due to inconsistent taps). The mean preferred tapping frequency was 1.55 Hz.
